# External morphometric and microscopic analysis of the reproductive system in *in-vitro* reared stingless bee queens, *Heterotrigona itama*, and their mating frequency

**DOI:** 10.1101/2024.06.12.598741

**Authors:** Kanyanat Wongsa, Orawan Duangphakdee, Pisit Poolprasert, Atsalek Rattanawannee

**Author notes:** Corresponding author (AR).

## Abstract

Stingless bees, prevalent in tropical and subtropical regions, are a tribe of eusocial bees that are crucial pollinators for economic crops and native plants, producing honey and pollen. However, colony expansion is limited by a shortage of queens for new colonies. Therefore, mass artificial rearing of virgin queens could address this in commercially managed meliponiculture. Furthermore, the in vitro rearing of queen stingless bees can improve meliponiculture management and conservation efforts. Herein, we explored the efficacy of in vitro queen rearing for *Heterotrigona itama* assessing the queen’s body size, reproductive organ size (ovary and spermatheca), acceptance rate into new, small colonies, and mating frequency. *H. itama* larvae developed into queens when fed with 120 µL–150 µL of larval food, resulting in *in vitro* queens having body sizes similar to those of naturally produced queens. Microscopic analysis revealed well-developed ovaries and spermathecae in *in vitro-*reared queens, unlike the smaller ovaries and the absence of spermathecae in the naturally produced workers. Acceptance of *in vitro-*reared queens was independent of worker age, and mating frequency was low but not significantly different from naturally produced queens. These findings could enhance stingless beekeeping practices and conservation efforts for the native stingless bee species.

## 1. Introduction

Stingless bees (Apidae: Meliponini) are eusocial bees found across tropical and subtropical regions [1–3]. They form perennial colonies comprising several hundreds to thousands of female workers [2]. These bees serve as highly effective pollinators for economic crops and native plants, playing a crucial role in biodiversity and agriculture [4–8]. Additionally, they are valued for their colony production, offering an abundant yield of honey and pollen [3, 9].

Meliponiculture (stingless beekeeping) has seen significant growth in Thailand, especially in rural villages, where it is promoted as an additional activity. The rich diversity of stingless bee species has fueled this revival. However, several Thai stingless beekeepers face challenges in multiplying the colonies with the original colonies, often due to the limited availability of virgin queens for successful propagation. Additionally, recognizing the right time [10] and developmental stage of the brood comb is crucial for multiplying stingless bee colonies. Therefore, this scarcity of virgin queens for establishing new colonies may greatly compromise management strategies for propagating stingless bee colonies [11].

Unlike honeybees of the genus *Apis*, stingless bees do not progressively feed larval food to their broods. Instead, they practice mass provisioning, preparing a mixture of gland secretions, honey, and pollen as larval food. This mixture is placed in a brood cell, and the queen lays an egg on top of it. Workers subsequently close the brood cells that remain open until adult stingless bees emerge [12, 13]. Most Meliponine species rear young queens in the largest brood cells, called royal cells, whereas workers and males are reared in smaller ones [14, 15]. Royal brood cells can contain up to eight times more larval food than worker broods in some stingless bee species [14]. Consequently, female larvae develop into queens when they receive the most larval food [15, 16].

Artificial queen-rearing techniques for stingless bees involve overfeeding 1–3-d-old female larvae *in vitro* [15–17], as caste determination is primarily based on the amount of larval food [18]. Female larvae destined to become queens receive more food in larger royal cells, whereas those becoming workers are reared in smaller brood cells [18]. *In vitro* rearing of virgin queens can address the low natural production of virgin queens in stingless bee species, leading to a rapid increase in new colonies [11, 19, 20].

At least 33 stingless bee species have been reported in Thailand [21–23]. Among them, *Heterotrigona itama* (Cockerell, 1918) is particularly effective in artificial wooden hive boxes. Beekeepers propagate *H. itama* colonies for sale, honey production, and renting for pollinating economic crops. These colonies can be sold for 3000–5000 THB (85–140 USD) each (AR: personal survey). Additionally, Thai *H. itama* honey sells for 1200–1500 THB (35–45 USD) per kilogram, which is approximately ten times the price of honey from Thai Apis mellifera and approximately three times that of honey from native Thai *Apis* species (*A. florea*, *A. dorsata*, and *A. cerana*) [3].

Hence, this study examined the impact of larval food quantity on the external morphology and reproductive organs of *in vitro*-reared *H. itama* queens. Furthermore, we determined the acceptance rate of these queens into small artificial queenless colonies and compared the mating frequency between *in vitro*-reared and naturally raised *H. itama* queens in commercial apiary conditions. These findings suggest an efficient technique similar to *Apis mellifera* practices (apiculture) for rapidly increasing *H. itama* colonies, enabling their availability for honey production and pollination. This approach improved meliponiculture management in Thailand and could help prevent overhunting of wild stingless bee colonies.

## 2. Materials and methods

### 2.1 Stingless bee and study sites

The experiments were conducted in the Department of Entomology, Kasetsart University, Bangkok, Thailand. Furthermore, the amount of larval food and acceptance of *in vitro*-reared queen into new colonies were determined at two commercial apiaries located in Yi-ngo district, Narathiwat province (06° 23′ 31′′ N; 101° 41′ 46′′ E) and Chulabhorn district, Nakhon Si Thammarat province (08° 01′ 06′′ N; 99° 48′ 29′′ E), southern Thailand. The stingless bee species used was *H. itama* (Cockerell, 1918). This species is a medium-sized stingless bee, with a worker body length of approximately 5.5−5.6 mm. Perennial colonies consist of approximately 8,000–10,000 worker bees and a single physogastric queen.

### 2.2 Ethics statement

Stingless beekeepers were prepared to employ this commercially significant species in this study, as they offered their approval in accordance with the statement of purpose and methods outlined in the consent form, agreeing to carry out the study on their own farms.

Farm owners then participated in colony quality evaluation and facilitated experimental arena preparation. Additionally, no endangered or protected species were included in the field survey. The sample number collected was kept to a minimum, and ethical treatment was applied according to the research standards. The Animal Experiment Committee of Kasetsart University, Thailand (approval no. ACKU66−AGR−015) approved all the animal-based experiments.

### 2.3 Amount of larval food

#### (a) Procedure to determine the amount of larval food

We assessed and compared the larval food amount in the queen brood cells (n = 82) to that in the worker brood cells (n = 290) of *H. itama*. These brood cells were collected from 16 unrelated parental colonies of two commercial apiaries in Narathiwat and Nakhon Si Thammarat provinces.

Ideal conditions for the colonies include numerous individual workers and strong single queens, as well as the absence of diseases and parasites.

Brood combs were collected in a manner that did not harm the sanity of the colonies. To avoid massive colony destruction, the recently built queen and worker brood cells were gently removed from the hive box using a knife and scalpel. Subsequently, the brood cells were kept in plastic containers (15 × 20 × 7 cm) in the dark to minimize potential microorganism exposure. The queen cells were separated from the edges of the brood combs to prevent excessive cell damage.

Calibrated microcapillary tubes with an auto-micropipette setup [16], with slight modifications, were used to meticulously collect and determine the larval feeding amounts from individual recently capped-brood cells, each containing a single egg [17]. Typically, newly constructed worker brood cells are dark brown, whereas as they progress into larvae, they lighten in color since the worker bees remove the cerumen [10].

#### (a) (b) Comparing the larval food amounts

We employed a one-sample Kolmogorov–Smirnov test to evaluate data normality to compare the larval food amounts between the queen and worker brood cells. After discovering that the data were non-parametric (Kolmogorov–Smirnov; Z = 8.627, *P*<0.001), we used a chi-squared test to compare the two groups (queens vs. workers). We first evaluated data normality using the one-sample Kolmogorov–Smirnov test to compare the larval food amount in the queen cells between the two meliponaries. As the data were also non-parametric (Kolmogorov–Smirnov; Z = 1.739, *P*=0.0047), we used the Mann–Whitney U test to compare between meliponaries. All data analyses were performed using IBM SPSS Statistics 22 [24].

### 2.4 Procedure involving *in vitro* queen rearing

#### (a) Harvesting larval food

This procedure aimed to collect adequate larval food for *in vitro* queen rearing. As previously mentioned, recently capped worker broods were removed from the hive boxes. To minimize colony disruption, only one or two layers of new worker brood cells were taken from each cell. Brood cells were subsequently opened using fine-tipped forceps. Before collecting the larval food, the eggs were removed using a sterile needle. Brood combs without eggs were transferred into a 50-ml centrifuge tube and subsequently squeezed to extract the larval food liquid. Each acrylic 96-well enzyme-linked immunosorbent assay (ELISA) plate was filled with the larval food liquid. In this study, we compared two larval food levels, 120 µL and 150 µL, to determine the quantity of the larval food required to produce queens equal in size to those naturally produced. This was based on the results of the present experiment, which determined the larval food amount in the queen brood cells (see Results), and the findings of Razali et al. [10]. Additionally, we added approximately 20% (making a total of 150 µL) of the same larval food amount to the normal queen cells [15] to ensure sufficient food for larval queen development.

#### (a) (b) Queen rearing procedure

In the laboratory, larval food of 120 µl (three plates; n = 288 wells) and 150 µl (three plates; n = 288 wells) were transferred into each well of the 96-well ELISA plates, providing sufficient space for each *H. itama* larva. First, the larval food was homogenized using a manual pipette (drawing up larval food and dispensing it several times) and was provided to the artificial cells. The new brood cells containing eggs were carefully opened, and the eggs were subsequently carefully removed from the brood cells using a sterilized beekeeping moving queen grafting tool (Fig. 1A) and transferred to each well (Fig. 1B). To retain their original orientation, all eggs were vertically placed on the larval food.

**Figure 1.**
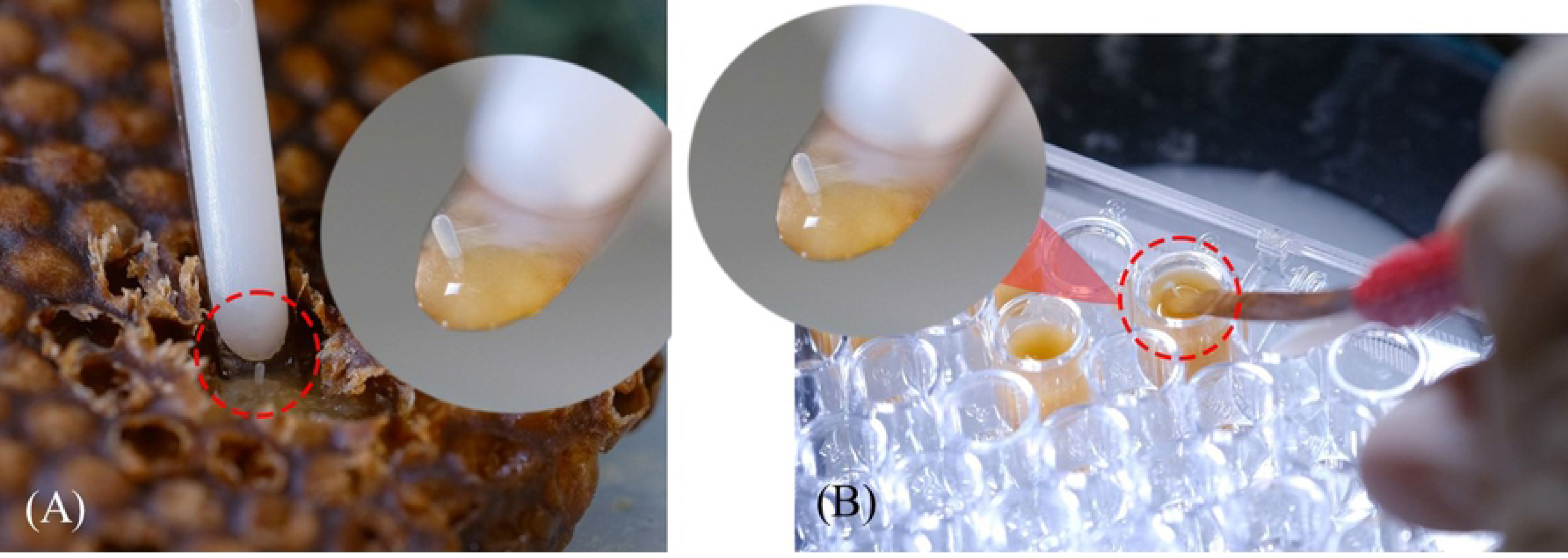
Representation of the egg-transferring phases of the *in vitro* rearing of the *Heterotrigona itama*. (A) The queen-laid egg was taken out carefully from the brood combs using a sterilized beekeeping moving queen grafting tool. (B) The queen-laid egg was placed in ELISA plate vertically as it is depos it edinnatural brood cell.

#### (a) (c) Incubation

After egg transfer, each queen-rearing plate was placed in hermetic plastic containers (12 cm × 25 cm × 6 cm) and housed in an incubator at 30°C (Binder SB incubator, Germany) with constant darkness (0 L: 24 D). Relative humidity in the plastic container was maintained between 70% and 100% throughout the larval growth phase using a sodium chloride (NaCl) saturated solution in distilled water as necessary (Table 1). Using Datalogger devices [(model: DHT22; sensor size: 22 mm × 28 mm × 5 mm, accuracy: humidity ±2% relative humidity (RH) (Max ±5% RH); temperature < ±0.5°C)], humidity data were obtained.

**Table 1.**
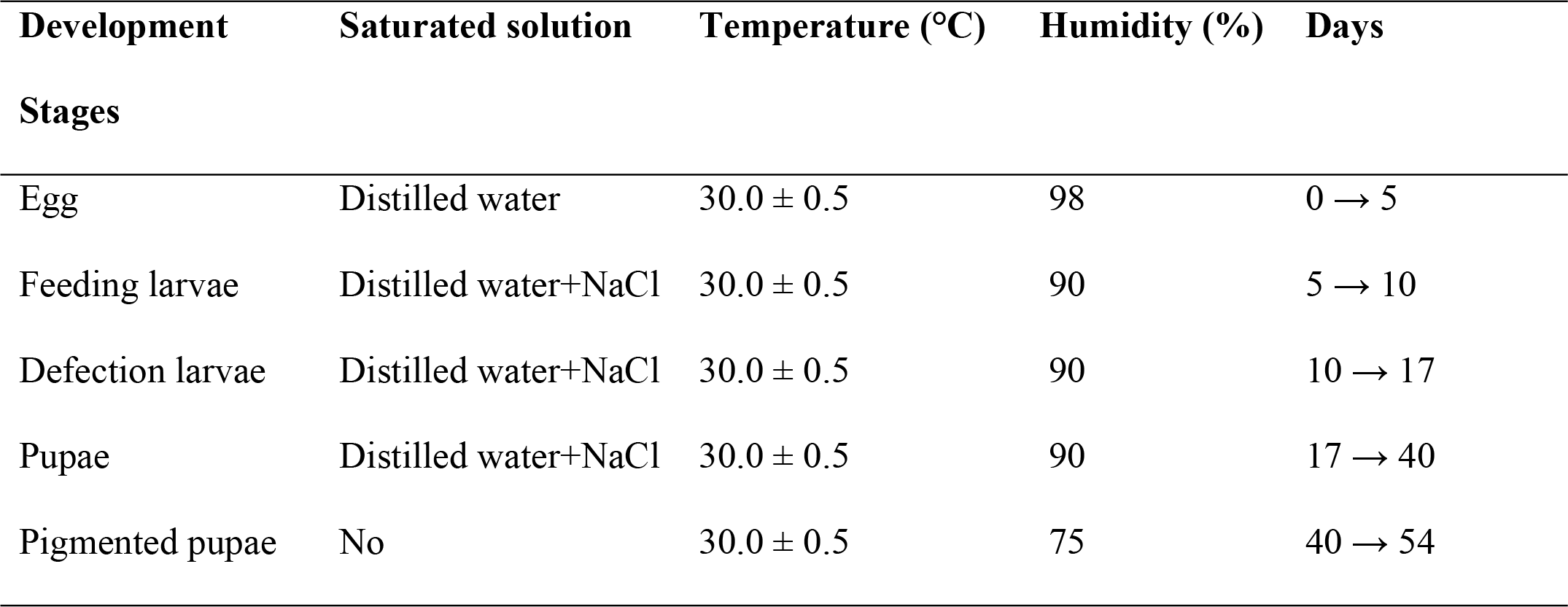
Overall features of the queen-rearing protocol of *Heterotrigona itama*

### 2.5 Microscopic analysis of the reproductive system

We conducted a comparative analysis of the ovaries of *in vitro* queens and newly emerged workers obtained from the *H. itama* natural colonies to explore virgin queen production under *in vitro* conditions. Workers were recruited randomly from the apiaries for this purpose. Fresh samples from *in vitro* queens (n = 11) and newly emerged workers (n = 20) were placed in petri dishes.

Subsequently, these samples were fixed in a phosphate-buffered saline (136.9 Mm NaCl, 8.1 Mm Na2HPO4, 2.7 Mm KH2PO4) with a pH of 7.1 to prevent the reproductive organs from drying out during the dissection process, following Razali and Razak’s protocol [10]. The dissection was performed using super-fine-tip forceps under an Olympus SZ51 stereomicroscope for precision.

Immediately following the dissection, images of the reproductive system were captured using a Leica EZ4W microscope.

### 2.6 Morphometric analysis

#### (a) Obtained natural virgin queen of H. itama

Thirty capped queen cells were collected from two apiaries located in the Nakhon Si Thammarat and Narathiwat provinces. Capped queen cells were kept separate from each other. Next, each queen cell was placed in an artificial cage and reintroduced into the colony from whence it originated. Adult virgin queens in the queen cages were collected upon emergence and preserved in 70% ethanol for morphometric analysis.

#### (a) (b) Morphometry measurement

All *in vitro* queens (n = 11), natural virgin queens (n = 22), and young adult workers (n = 30) were dissected using fine-tipped forceps under a stereomicroscope. This method included the antenna, mandible, head (with compound eyes), right forewing, right hindwing, right hindleg (with the femur, tibia, and basitarsus), 4^th^ and 5^th^ tergites, and 4^th^ and 5^th^ sternites. All body parts were mounted onto microscope slides and photographed using a digital camera attached to a stereomicroscope (Leica EZ4 W/E; Leica). Leica LAS EZ software (Leica, Wetzlar, Germany) was used for measuring all morphometric traits according to previously published methods [25–28]. Herein, 34 morphometric characteristics of the bee samples were measured and analyzed (Fig. S1).

#### (a) (c) Morphometry analysis

Means and standard errors (SE) of all characteristics were calculated. The *G*-test, which is included in IBM SPSS Statistics 22, was used to examine the skewness of the character sizes [24]. Discriminant analysis was used to discriminate between the differences in external characteristics among the *in vitro* virgin queens, natural virgin queens, and young adult workers. Samples were grouped into three categories: *in vitro* virgin queens, natural virgin queens, and young adult workers to compare different sizes. The characters’ centroid size (CS) was calculated as a measure of the overall size of the stingless bee specimens [29] and was used to assess whether bees from various groups differed in size. The CS of all characters among the three *H. itama* groups was compared using discriminant analysis implemented in IBM SPSS Statistics 22 [24].

### 2.7 *In vitro* queen acceptance

#### (a) Introducing the in vitro queen into the new colony

We investigated the acceptance of queens reared *in vitro* into new artificial queenless colonies. First, we produced two sets of artificial small queenless colonies (n = 27) by introducing approximately 500 workers of various ages (artificial queenless colonies a (AQCa): newly emergent workers only; artificial queenless colonies b (AQCb): newly emergent and old workers) collected from strong queen-right colonies. Second, we introduced two brood combs containing approximately 200 larvae close to emergence. Finally, approximately 100 g of pollen and 200 g of honey pots were placed in the corners of the hive box. After 2–3 d, approximately one-week-old queens reared *in vitro* were introduced into queen cages (Video S1) and subsequently placed inside artificial queenless colonies (AQCA = 14 colonies and AQCB = 13 colonies). Their behavior toward the workers was recorded. If the *in vitro* queens remained alive for at least 7 d after introduction, they were declared as being accepted. Additionally, we deemed the *in vitro* rearing of the queens as successful when the queens vibrated their wings, were not attacked by workers, and exhibited trophallaxis with other bees [16]. All experimental procedures were conducted at the Nakhon Si Thammarat apiary.

#### (a) (b) Analyzing the acceptance of in vitro queens

We evaluated the likelihood of the successful *in vitro* adoption of the queens into artificial queenless colonies using an odds ratio test with a 95% confidence interval (CI). The acceptance rate of the *in vitro* queens was compared between two types of artificial queenless colonies containing (a) only non-pigmented workers and (b) non-pigmented and old workers using Pearson’s chi-squared test (n = 9999; Monte Carlo simulation). All data analyses were performed using IBM SPSS Statistics 22 [24].

### 2.8 Analyzing the mating frequency of the *in vitro* queen

#### (a) Worker sample collection

Ten young adult workers were collected from three-month-old colonies containing *in vitro* queens and preserved in 95% (v/v) ethanol to evaluate the number of fathering genotypes of *in vitro* queen-headed colonies. All young adult worker samples were directly collected from the brood combs inside each colony. The samples were stored at −20°C until further use for DNA extraction. Eight *in vitro* queen-heading colonies were used in this study.

#### (b) DNA extraction, polymerase chain reaction (PCR) condition, and genotyping

Genomic DNA was extracted from the right hind leg of individual worker bee using a 5% (w/v) Chelex solution (Chelex®100; BIO-RAD), as followed by Walsh, Metzger (30) with slight modifications. Three microsatellite target sequences were amplified via PCR using fluorescent-labeled primers. The microsatellite loci used were TC4.287, TC7.13, and TC3.155 [31]. Table S1 lists the primer details. The PCRs were conducted using a T100™ thermal cycler (BIO-RAD) containing 1× Multiplex PCR Master Mix (Green HotStart PCR Master Mix, Biotechrabbit), 1 µL of each primer (20 µmol/L), and 1 µL of genomic DNA template, and distilled water up to a total volume of 10 μL. All microsatellite loci were amplified using a standard PCR program of 94°C for 5 min, followed by 35 cycles of 94°C, 56°C, and 72°C for 30 s each, and finally 72°C for 10 min. Samples without DNA were included in all the plates as negative controls. The PCR products were subsequently sent to Macrogen (Macrogen, Korea) for fragment analysis. The resulting data files were analyzed for allele size determination using GENEMAPPER (Applied Biosystems).

**Table S1** Microsatellite loci used.

#### (c) Reconstructing the queen genotype and identifying patrilines

The queen heading genotype for each colony was inferred from the worker genotypes [32–34]. Workers were deemed drifting individuals if they did not carry the queen allele [34]. After determining the queen’s genotype, the fathering drone genotype was determined for each worker by subtraction [32, 34, 35].

#### (d) Analyzing the mating frequency

The effective mating frequency (*me*) within each colony, with correction for a finite sample size, was calculated according to the method followed by Tarpy and Nielsen (36). To determine intracolonial relatedness, the average relatedness, *r*, weighted according to the relative proportions of each subfamily and corrected for finite sample size, was calculated for each colony according to the method followed by Oldroyd and Moran (37). Furthermore, the observed paternity frequency (*k)* was calculated for a common number of workers scored according to the method followed by Franck, Koeniger (38) to make a valid comparison of the mating frequency between natural and *in vitro*- reared queens.

## 3. Results

### 3.1 Amount of larval food in the queen and worker brood cells

This experiment aimed to determine the larval food amount required for the growth of *H. itama* queens and their workers under natural colony conditions. The larval food amount deposited in queen brood cells (119.53 ± 2.42 µL) was significantly higher than those found in the worker brood cells (18.22 ± 1.67 µL) in the normal queen-right colonies (ꭓ^2^ = 434.77, *df* = 90, *P* < 0.001) (Fig. 2). The differences in the larval food amount deposited in the queen brood cells between apiaries (Mann– Whitney U test: *U* = 823.5, *P* = 0.878) and among colonies were insignificant (Kruskal–Wallis Test: ꭓ^2^ = 19.43, *df* = 15, *P* = 0.195). Notably, the larval food deposited in the queen brood cells was approximately seven times higher than that deposited in the worker brood cells under natural conditions.

**Figure 2.**
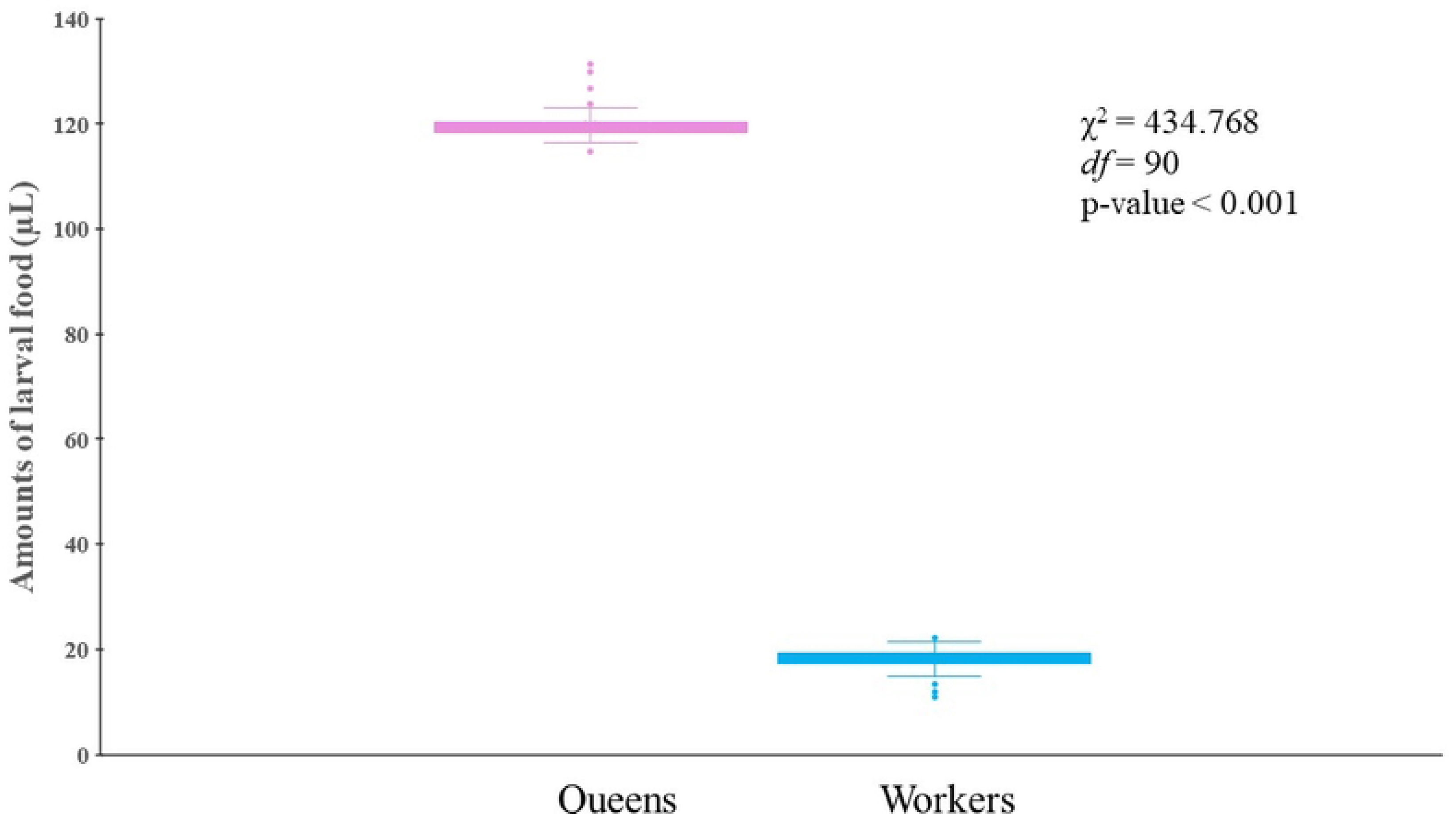
Comparison between the amounts of larval food found in natural brood cells of queens and workers of *Heterotrigona itama*. The boxplot displays median values, 1^st^ and 3^rd^ quartiles, and the upper and lower lines indicating the maximum and minimum values.

### 3.2 *In vitro* larval development

Depending on the larval developmental stage, the average temperature and humidity in an incubator filled with distilled water and NaCl were maintained (Table 1). In this experiment, the eggs took approximately 4-5 d to develop into larval forms. After hatching, the larvae began feeding on the food provided on the acrylic plates. Subsequently, the larvae turned into pupae after approximately 17 d (Table 1 and Fig. 3). Larval queens reared *in vitro* developed for approximately 54 d before emerging as adults in this study.

**Figure 3.**
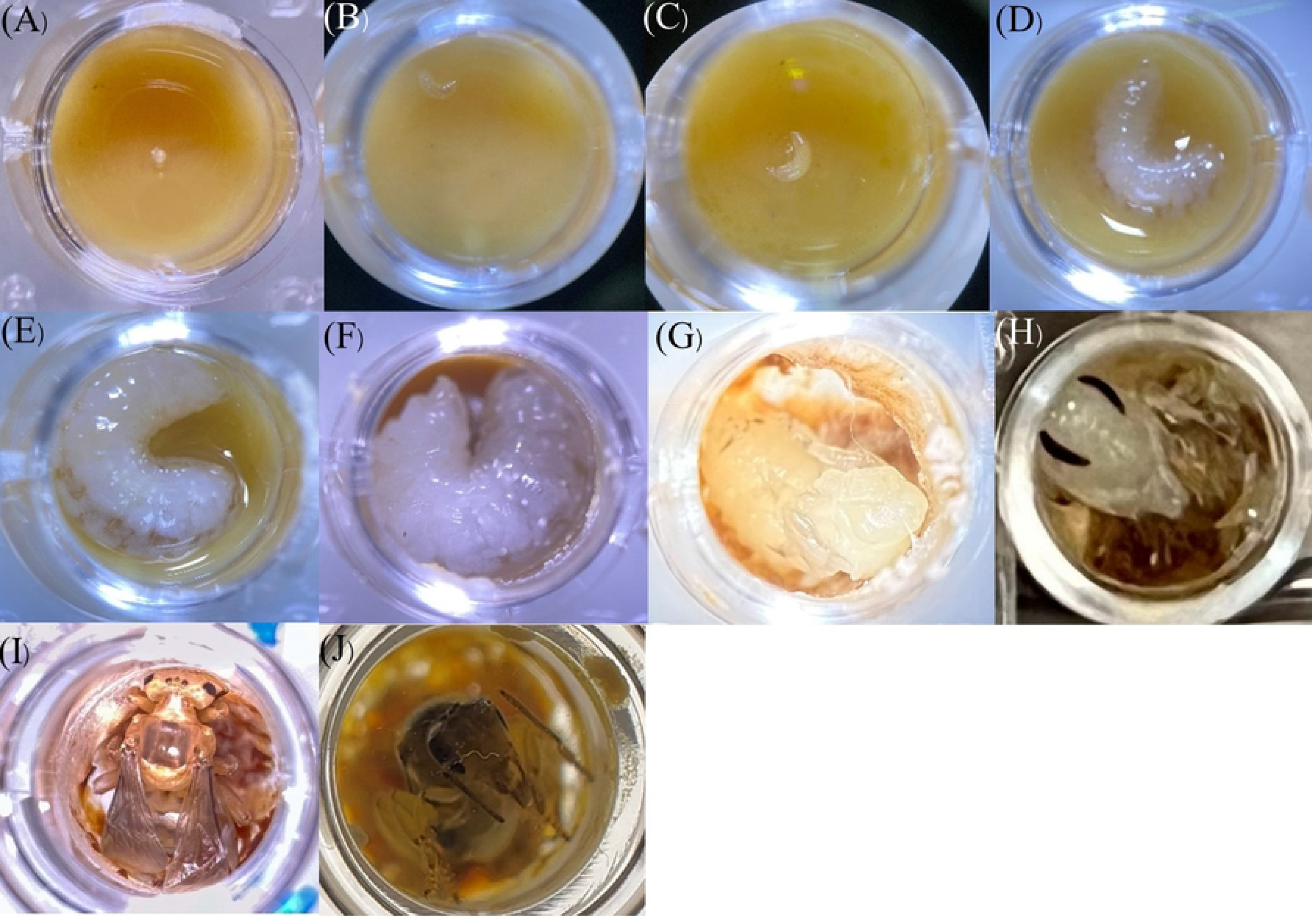
Observation of the *in vitro* development stages of *Heterotrigona itama* queen larvae under a stereo microscope with 1.5X magnification, spanning from (A) eggs, (B) to (E) feeding larvae, (F) prepupae, (G) white-eyed pupae, (H) brown-eyed and white-bodied pupae, (I) pigmented pupae, and (J) adult queen.

### 3.3 Worker and queen emergence in response to the amount of larval food

This study yielded 173 (60.07 ± 5.92%) and 186 (64.58 ± 3.76%) *in vitro H. itama* queens from 120 µL and 150 µL of larval food, respectively. The two larval food levels provided in response to queen emergence did not significantly differ (*t* = 1.115, *df* = 4, *P* = 0.327). The eggs treated with the highest larval food amount (150 µL) resulted in 64.58 ± 3.76% queen and 6.94 ± 2.62% worker emergence (Fig. 4, and Video S2 and S3). The results of providing 120 µL of food revealed a lower queen emergence percentage (60.07 ± 5.92%); however, this rate was not significantly different from that obtained when 150 µL of food was provided. However, both food levels resulted in a high dead larval percentage (Fig. 4).

**Figure 4.**
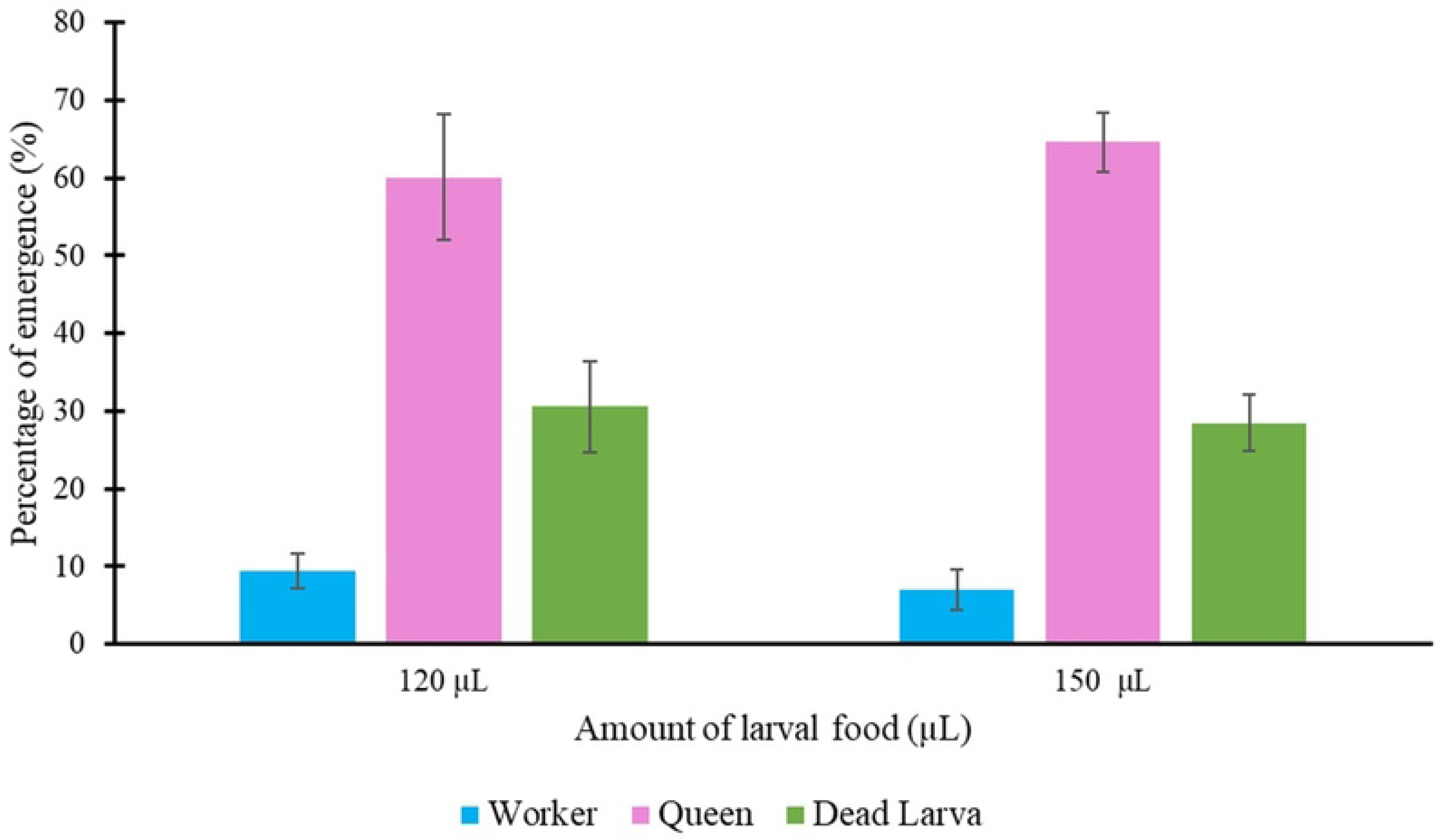
The percentage of worker and queen emergence of *Heterotrigona itama* in respond to larval food amounto f120 µ L and 150 µ L from the *in - vitro* rearing experiment.

### 3.4 Microscopy analysis of the reproductive system

We compared the reproductive systems of the *in vitro* queens with those of naturally produced worker stingless bees to verify whether the *in vitro*-produced adult is a queen. Ovary size and the presence of a spermatheca were used to differentiate between queens and workers, conclusively confirming the status of an emerging adult as a queen. *In vitro* queens from the 120 and 150 µL treatments were larger than workers (Fig. 5), with a considerable abdomen and well-developed reproductive system. They possessed a large ovary with numerous ovarian filaments (ovarioles) and spermathecae (Fig. 6), which increased the likelihood of them laying eggs.

**Figure 5.**
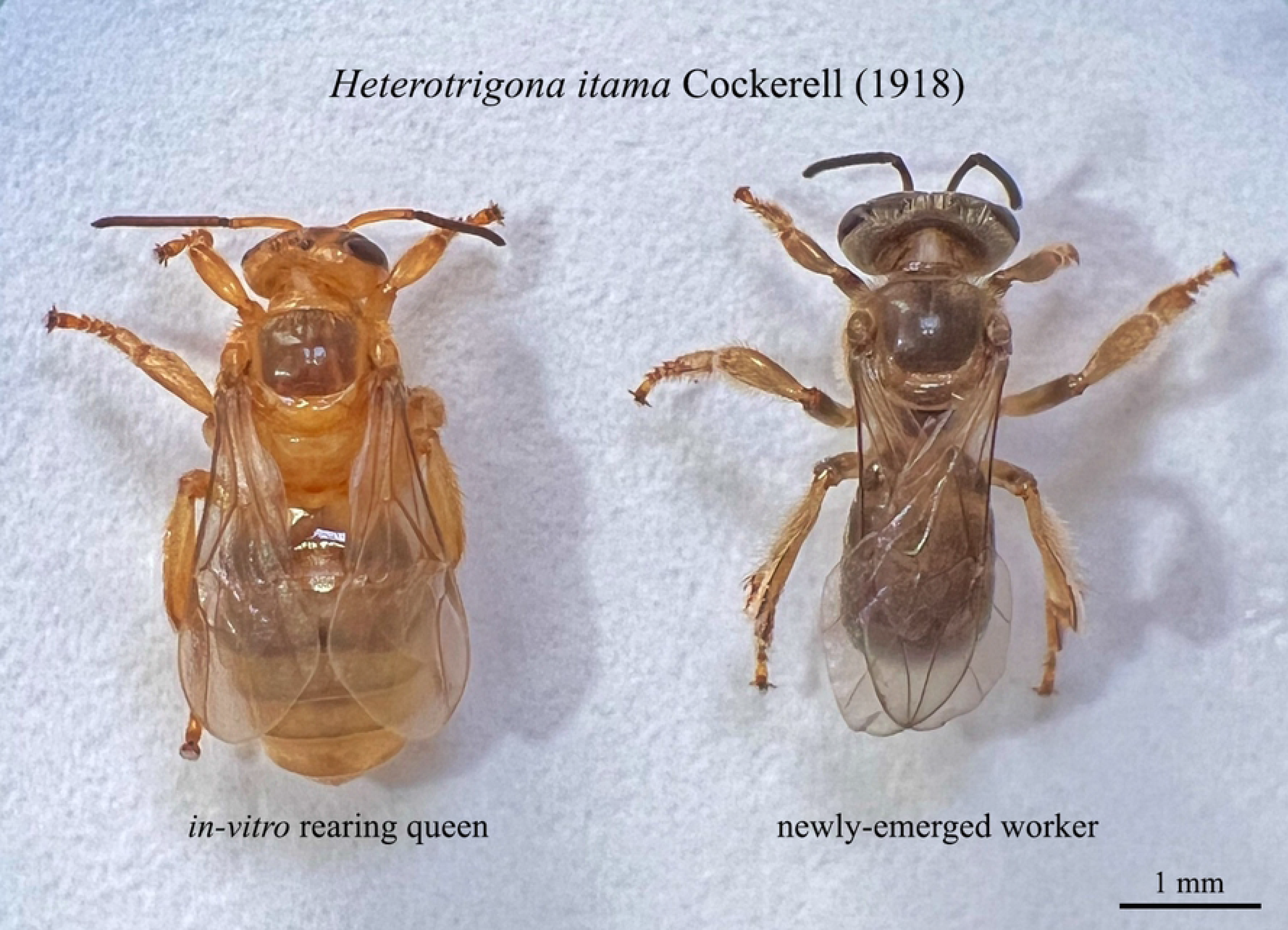
Newly emerged in-vitro rearing queen and worker and of *Heterotrigona itama*.

**Figure 6.**
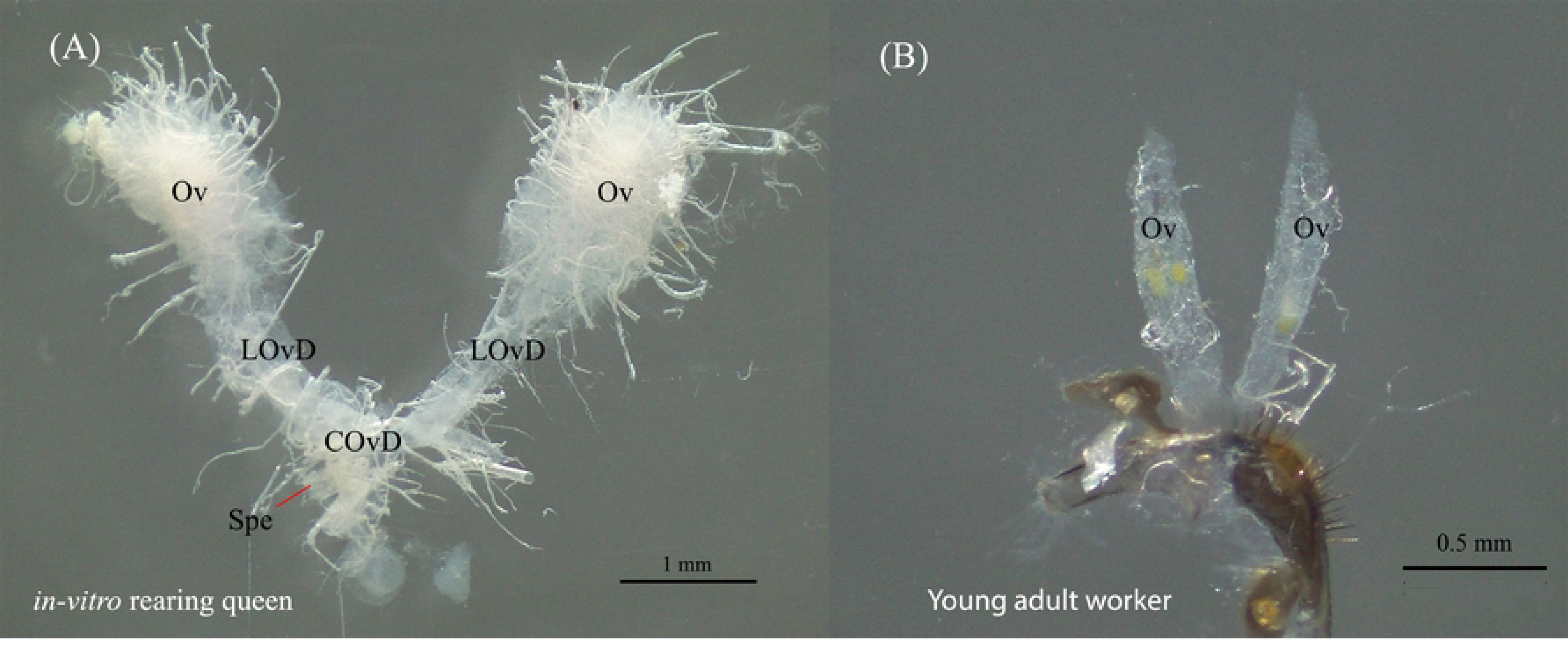
Microscopy images of gonadal development in *Heterotrigona itama*, displaying (A) the freshly dissected reproductive system of an *in-vitro* queen with a well-developed ovarioles structure, and (B) a dissection of female workers, highlighting significant differences in ovaries and ovariole size compared to the *in-vitro* queen. The abreviations are ovary (Ov), lateral oviduct (LOvD), commono viduct (COvD), and spermateca (Spe).

Contrastingly, *H. itama* workers had ovaries but no spermathecae and their ovary size was smaller with fewer ovarioles than that of the *in vitro* queens (Fig. 6).

### 3.5 Morphometric analysis

We assessed and compared 34 morphometric characteristics among the *in vitro* queens, natural virgin queens, and adult workers (Table S2). Discriminant analysis showed no significant differences between the *in vitro* and virgin queens produced under natural conditions; however, they were clearly separated from the workers (Fig. 7). Morphometric analysis demonstrated no significant differences in size between the *in vitro*-reared queens and natural virgin queens (*p* > 0.05), except for the head length (*p* = 0.019), antenna length (*p* = 0.017), length of tergite 4 (*p* = 0.023), and width of the upper edge of tergite 4 (*p* = 0.006), where the *in vitro* queens were found to be smaller than the natural virgin queens (Table S3). Furthermore, queens under both conditions were considerably larger than the workers, except for the number of teeth (*p* > 0.05) and flagellum segments (*p* > 0.05) (Table S3).

**Figure 7.**
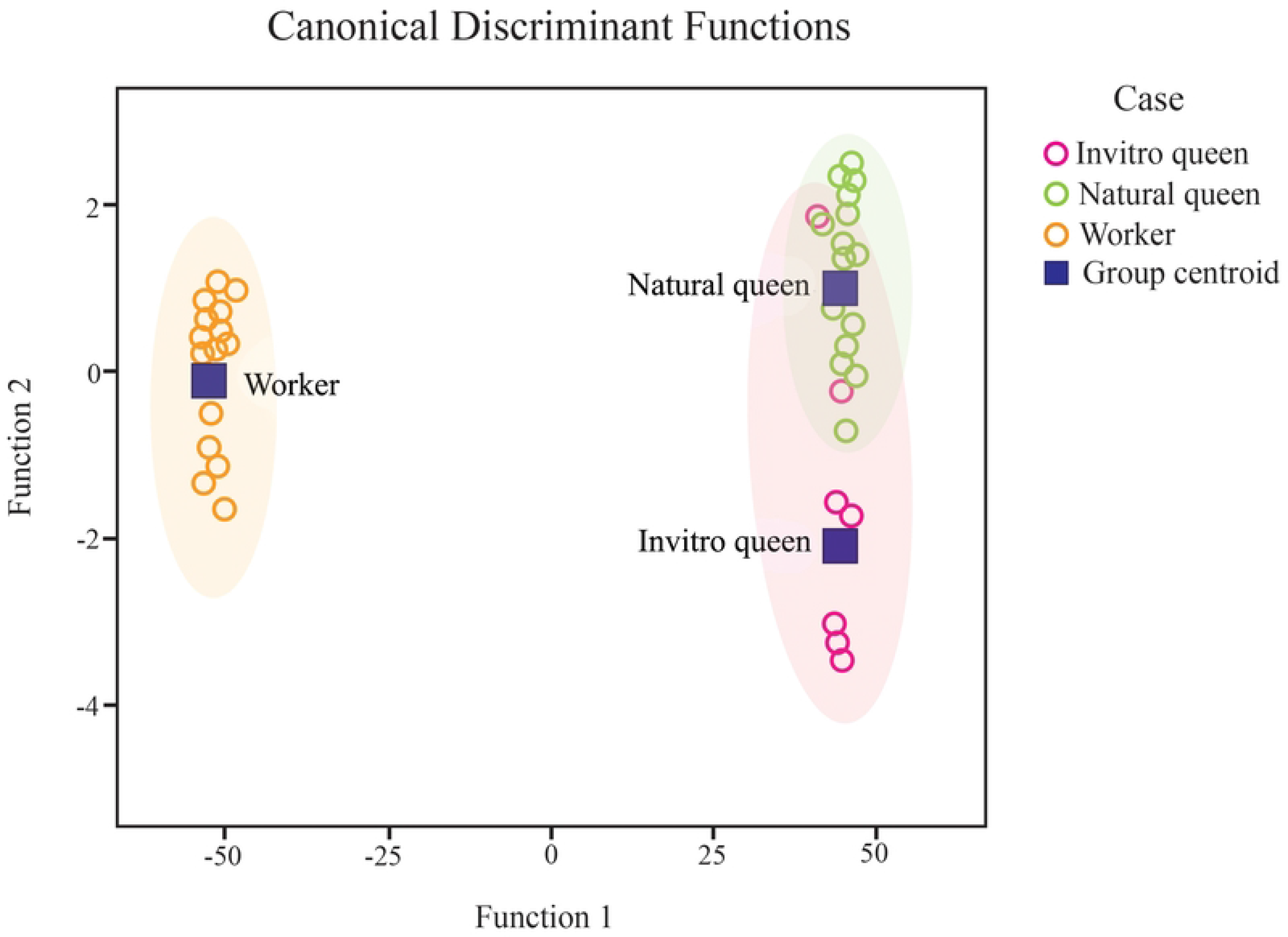
Linear discriminant analysis was conducted on the 34 morphometric characters measured in *in vitro*-reared queens, natural virgin queens, and workers of *Heterotrigona itama*.

Interestingly, workers exhibited markedly greater head width, compound eye length and width, mandible length, hind tibia length and width, hind basitarsus length, forewing length and width, marginal cell length, 1^st^ submarginal cell length, and hind wing length and width when compared to the *in vitro* and natural virgin queens (Table S3).

### 3.6 Acceptance rate of the *in vitro*-reared virgin queens

Over 70% of the *in vitro*-reared queens were accepted after being introduced to the artificial queenless colonies that consisted of young workers only (AQCa). Contrastingly, a lower acceptance rate (approximately 46%) of the *in vitro*-reared queens was found in the queenless colonies that consisted of young and old workers (AQCb) (Fig. 8). The odds ratio (OR) test demonstrated that the *in vitro*-reared queens had a high acceptance rate in small artificial colonies that contained only young worker bees (OR = 2.917; 95 % CI 0.594‒14.). The difference in the likelihood of rejecting the *in vitro*-reared queens between older workers from small queenless colonies was insignificant (ꭓ2 = 1.784, *df* = 1, *P* = 0.182).

**Figure 8.**
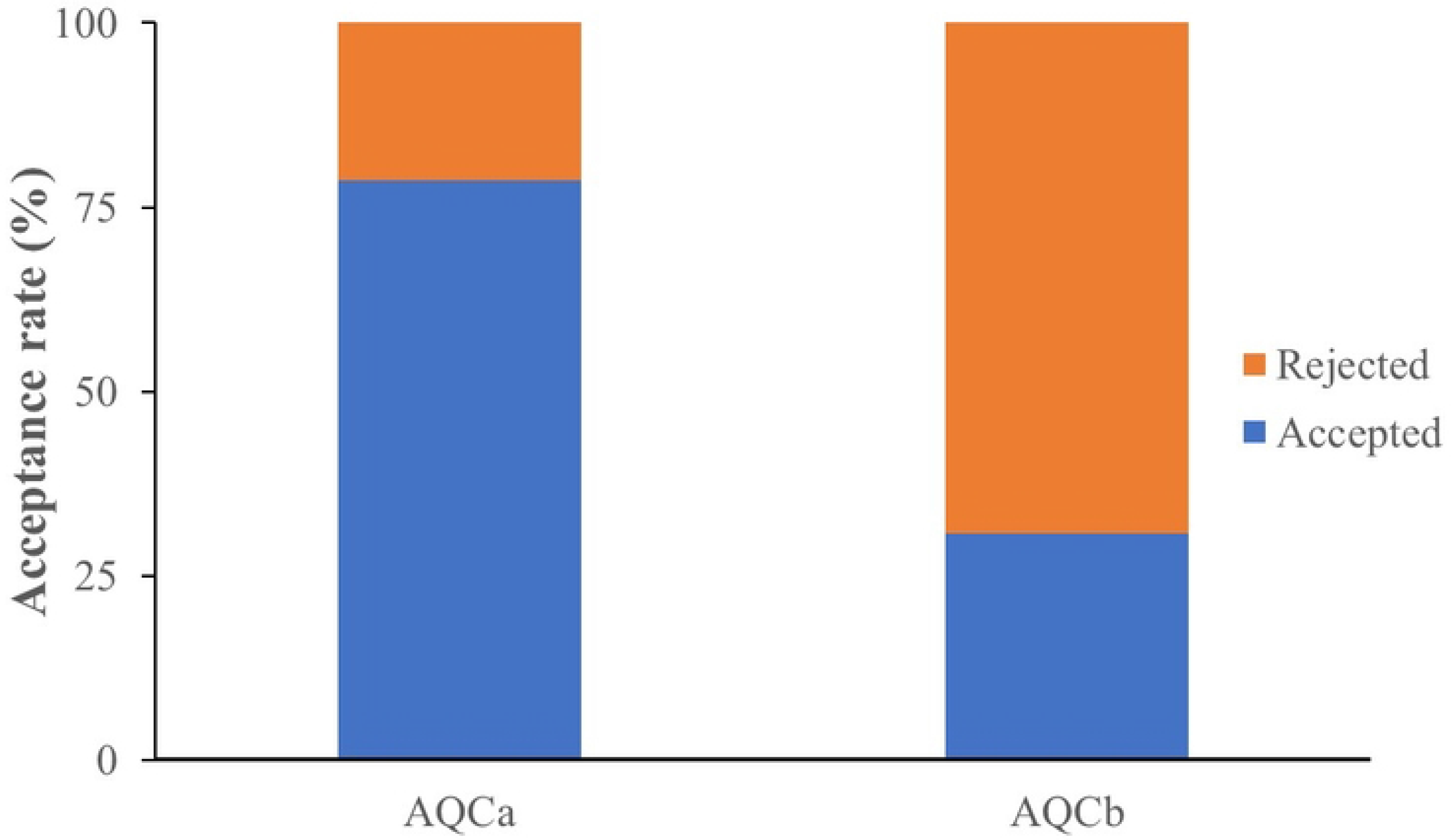
Acceptance rate of *Heterotrigona itama* queens reared *in vitro* inside artificial queenless colonies containing young workers only (AQCa) or young and old workers (AQCb).

### 3.7 Paternity frequency and intracolonial variation

The observed mating frequency, *k* (*t* =0.389, *df* = 13, *P* = 0.704), effective mating frequency,

*m_e_* (*t* =0.502, *df* = 13, *P* = 0.624), mating frequency corrected for sample size, *k_9_* (*t* =0.810, *df* = 13, *P* = 0.442), and intra*-*colonial relatedness, *r* (*t* =0.295, *df* = 13, *P* = 0.772) were not significantly different between the natural and *in vitro*-reared queens. Low effective paternity frequency values (2.086 ± 0.296) were observed in the *H. itama* queens, ranging from 1.00 to 3.37 (Table 2).

**Table 2.**
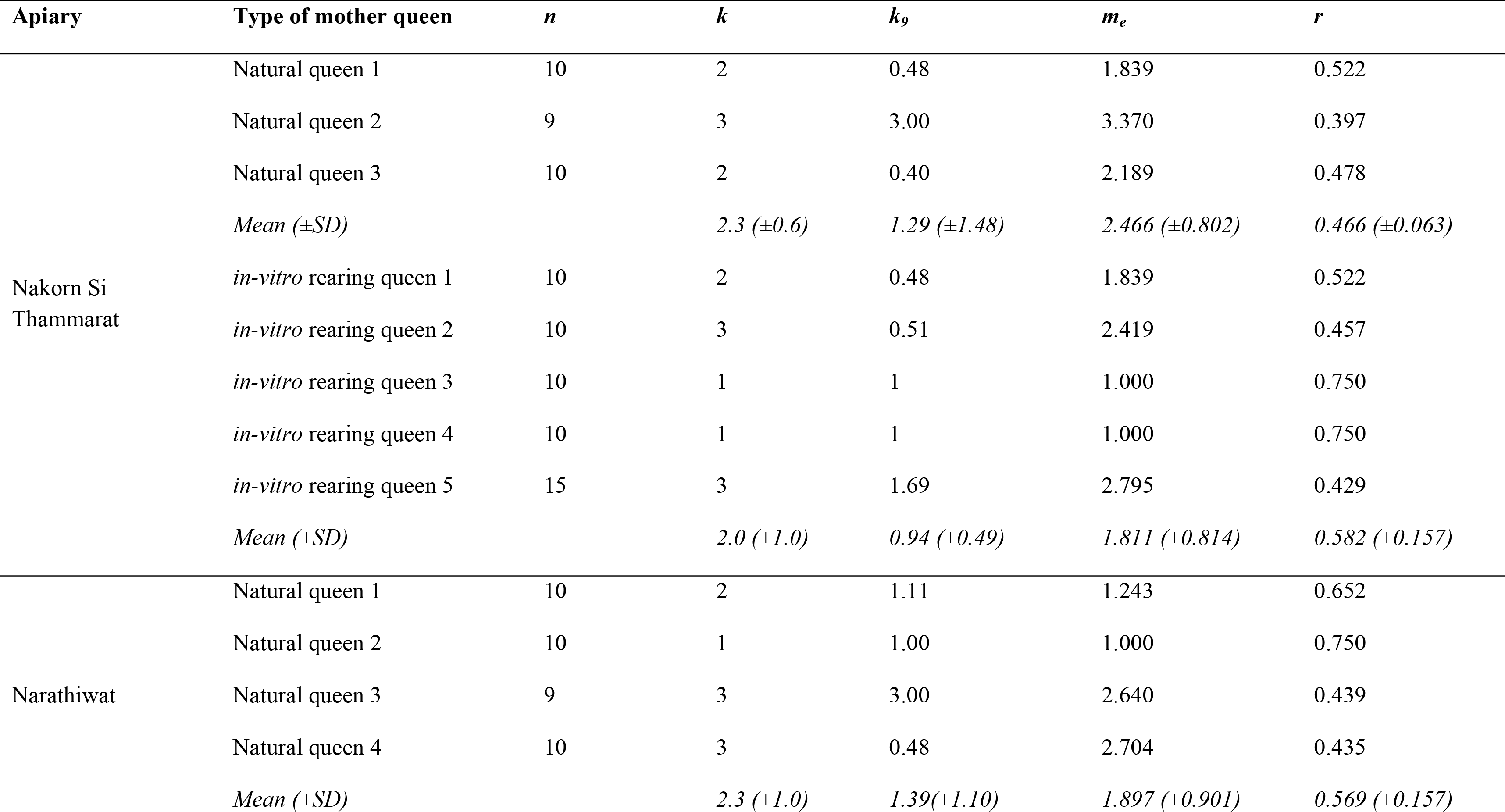
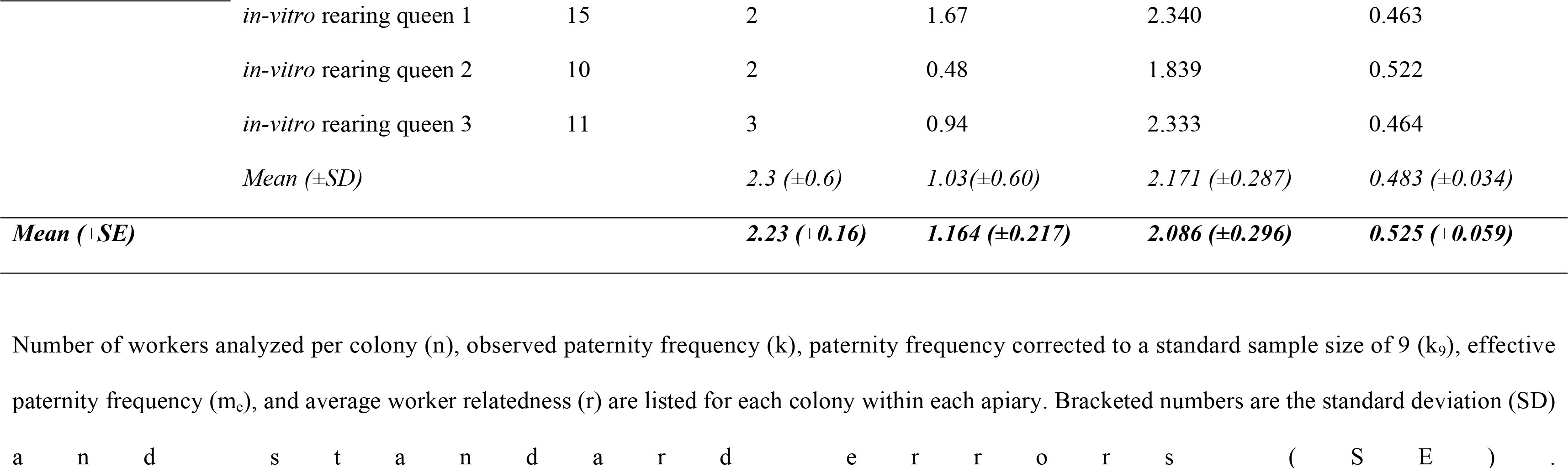
Paternity frequency in *Heterotrigona itama* colonies from two commercial apiaries in Thailand.

Additionally, the differences in the effective paternity frequency (*m_e_*) were insignificant and we deduced the paternity frequency (*k_9_*) between the Nakhon Si Thamrat and Narathiwat apiaries (*t* =0.108, *df* = 13, *P* = 0.916 and *t* =0.371, *df* = 13, *P* = 0.716, respectively). No significant difference was observed in the *m_e_* and *k_9_* that were used to compare the natural and *in vitro*-reared queens within apiaries (Nakhon Si Thammarat; *m_e_*: *t* =1.107, *df* = 6, *P* = 0.311, and k_9_: *t* =0.519, *df* = 6, *P* = 0.623; Narathiwat; *m_e_*: *t* =0.497, *df* = 5, *P* = 0.640, and k_9_: *t* =0.515, *df* = 5, *P* = 0.629).

We detected no drifting workers between or among the colonies within each apiary (Table 2). This may be attributed to the worker samples being directly collected from the brood clusters inside each colony [33, 34, 39].

## 4. Discussion

In this study, rearing *H. itama* queen larvae *in vitro* with 120 µL and 150 µL of food resulted in queens with similar body sizes to natural virgin queens after 54 d (Fig. 4 and 7; Tables S1 and S2). Previous research on *in vitro* stingless bee queen rearing has noted the challenge of achieving naturally sized artificial queens. Hartfelder and Engels (13)] successfully produced numerous *in vitro*- reared queens of *Scaptotrigona postica* but found them smaller than naturally produced queens.

Baptistella, Souza (17) observed that *in vitro*-reared *Frieseomelitta varia* queens were smaller than those naturally produced. However, Menezes, Vollet-Neto (15) reported that adding approximately 10% more larval food than the average found in *Scaptotrigona depilis* natural royal cells resulted in *in vitro*-reared queens with the same body size as natural queens. Queen body size is crucial because smaller queens lay fewer eggs than larger queens in some stingless bee species [40, 41].

*In vitro*-reared queens fed with 120 µL and 150 µL of food exhibited larger abdomens, well-developed ovaries with several ovarioles, and a spermatheca, enhancing the chances of successful mating and egg laying (Fig. 5 and 6). However, we observed the highest rate of queen larval mortality between the last larval stage and the beginning of the pupation phase. Similar findings were noted in *Plebeia droryana*, with the highest loss of *in vitro*-reared queen larvae occurring between the last larval stage and prepupal phase [16]. Consequently, future research may need to focus on improvements such as humidity and temperature adjustments during this critical period.

Our findings revealed no significant differences in queen emergence between the two larval food levels (Fig. 4). However, the reproductive organs of *in vitro*-reared queens fed with 120 µL and 150 µL larval food substantially differed from those of naturally produced workers (Fig. 6). Workers had smaller ovaries with fewer ovarioles and spermathecae. These results clearly highlight the distinct differences in the female reproductive systems between in vitro-reared queens and workers, attributed to the different larval food quantities provided [15, 18]. Similarly, Razali, Razak (10)] reported consistent findings, demonstrating that *in vitro*-reared *H. itama* queens fed with 150 µL larval food exhibited substantially larger ovaries compared to those of natural workers.

The acceptance of *in vitro*-reared *H. itama* queens was unaffected by the age of the workers. In this stingless bee species, the age of workers did not markedly impact their acceptance of *in vitro-* reared queens. In contrast to *P. droryana*, Fernando dos Santos, de Souza dos Santos (16) found that small queenless colonies with older workers had higher rejection rates for *in vitro*-reared queens. This study observed no effect of old workers on acceptance rates, likely because virgin queens were released using a queen cage, reducing direct contact between them and workers minimizing the risk of old workers killing the virgin queens. In practice, young and old workers, as well as young and old brood combs should be used to establish new colonies, along with honey and pollen pots.

The mating ability of *in vitro*-reared queens of stingless bees was first reported by Camargo (42), who found that only one out of five *in vitro*-reared *S. postica* queens mated and successfully laid eggs. Hartfelder and Engels (13) successfully produced a considerable number of *in vitro*-reared *S. postica* queens capable of mating and laying eggs. However, this study observed differences in the shapes and sizes of reproductive organs between *in vitro*-reared and naturally produced queens.

Menezes, Vollet-Neto (15)] demonstrated that *S. depilis* could mate and lay fertile eggs. In our study, we observed a low but not significantly different mating frequency between successfully mated *in vitro*-reared queens and naturally produced *H. itama* queens (Table 2). *In vitro* rearing of *H. itama* queens emerges as an efficient method for producing numerous virgin queens with a high potential for adequate colony propagation.

To date, several stingless bee species, including *S. postica* [13], *S. depilis* [15], *Plebeia droryana* [16], *Nannnotrigona testaceicornis* [43], *Tetragonisca angustula* [44], *Frieseomelitta varia* [17], and *H. itama* [10], have been successfully reared using *in vitro* queen rearing techniques. While protocols varied slightly, the fundamental techniques employed in these studies were similar to those used in the present study.

*In vitro* queen rearing of stingless bees offers an efficient method for obtaining numerous valuable virgin queens to multiply colonies. When optimal conditions, including consistent body size, shape, and reproductive organ size, as well as a mating frequency akin to natural queens, producing several hundred virgin queens from a single mother colony in a short timeframe is possible. Thus, this method holds promise from colony propagation and genetic improvement efforts [15], akin to practices in apiculture [45]. Meliponiculture (stingless beekeeping), a significant economic activity, has seen a surge in honey production and crop pollination in tropical and subtropical areas.

Consequently, further advancements in breeding techniques and colony management practices are imperative [15, 19].

## 5. Conclusion

This study demonstrates that depositing 120 µL and 150 µL of larval food *Heterotrigona itama* royal cells led to the development of queens within 54 d. *In vitro* queen generation using these food quantities produced queens resembling natural queens but larger than natural workers, well-developed ovaries, and spermathecae. We observed substantial mortality, especially between the last larval stage and the start of the pupation phase. Furthermore, the age of the workers did not affect the acceptance rate of the *in vitro*-reared queens. Regarding mating ability, we observed a modest but non-significant frequency between successfully mated in vitro-reared queens and naturally produced queens. This study aimed to enhance colony multiplication over a short period for modern economic meliponiculture activities. This technique potentially increases the number of new colonies and the economic value for stingless beekeepers.

## Conflict of interest

The authors have no conflict of interest.

## Acknowledgments

We thank the many stingless beekeepers who participated and made their hives available for the experiments. This study was supported by grant from the National Research Council of Thailand (NRCT) and Kasetsart University (Grant No. N42A650288), and the Kasetsart University Research and Development Institute (KURDI) (Grant No. FF(KU) 52.67)

---

Table S1 Microsatellite loci used

Table S2 Morphometric characters measurements were taken for in vitro-reared queens (n=11), natural virgin queens (n=22), and workers (n=30) of *Heterotrigona itama* from Thailand.

Table S3 Multiple comparisons of 34 morphometric characters were conducted among *in vitro*-reared queens, natural virgin queens, and workers of *Heterotrigona itama*. The character abbreviations correspond to Table S2 and Figure S1. The asterisks indicate statistically significant differences.

Figure S1 Thirty-four morphometric characters of *in-vitro* queens, natural virgin queens, and young adult workers of *Heterotrigona itama* were examined, including (A) head, (B) antenna, (C) mandible, (D) forewing and hindwing, (E) hind femur and hind tibia, (F) 4th and 5th tergites, (G) hind basitarsus, (H) 4th sternite, and (I) 5th sternite.

**Video S1** One-week-old queen reared *in vitro* of *Heterotrigona itama* was introduced into queen cages and then subsequently placed inside artificial queenless colony.

**Video S2** The *in vitro*-reared queen of *Heterotrigona itama* is emerging from artificial brood cell.

**Video S3** The *in vitro*-reared worker of *Heterotrigona itama* is emerging from artificial brood cell.

## Notes

### Competing Interest Statement

The authors have declared no competing interest.

